# Harnessing cGAS-STING signaling to counteract the genotoxic-immune nexus in tauopathy

**DOI:** 10.1101/2025.09.27.678980

**Authors:** Himanshi Singh, Sazzad Khan, Jianfeng Xiao, Nicole Nguyen, Asmita Das, Daniel Johnson, Francesca Fang-Liao, Sally A. Frautschy, Michael P. McDonald, Tayebeh Pourmotabbed, Mohammad Moshahid Khan

**Affiliations:** Department of Neurology, College of Medicine, University of Tennessee Health Science Center, Memphis, TN, 38163, USA; Department of Microbiology, Immunology and Biochemistry, University of Tennessee Health Science Center, Memphis, TN 38163, USA; Department of Biotechnology, Delhi Technological University, Shahbad Daulatpur, Delhi 110042, India; Molecular Bioinformatics Core, Translational Science Research Building, Memphis, TN 38163; Neuroscience Institute, University of Tennessee Health Science Center, Memphis, TN, USA; Departments of Neurology and Medicine, University of California, Los Angeles Veterans Administration, Geriatric Research; Department of Anatomy & Neurobiology, College of Medicine, University of Tennessee Health Science Center, Memphis, TN, USA; Center for Muscle, Metabolism and Neuropathology, Division of Regenerative and Rehabilitation Sciences and Department of Physical Therapy, College of Health Professions, University of Tennessee Health Science Center, Memphis, TN, USA

**Keywords:** DNA damage and repair, cGAS-STING, immune response, tauopathy, senescence, cognitive function, neuroinflammation

## Abstract

Tauopathies are progressive neurodegenerative disorders characterized by aberrant tau aggregation, cognitive decline, and persistent neuroinflammation, yet the mechanisms driving neuroinflammation and disease progression remain incompletely understood. Here, utilizing human postmortem AD brains and a mouse model of tauopathy, we report that genotoxic stress-induced cyclic GMP-AMP synthase (cGAS)-stimulator of interferon genes (STING) immune pathway form a self-amplifying loop that fuels neuropathology and cognitive deficits. Targeted disruption of this cycle through either genetic deletion of cGAS or pharmacological inhibition of STING restores immune homeostasis and attenuates tau pathology and cognitive deficits. Our results showed a significant accumulation of DNA double-strand breaks (DDSBs) and impaired DNA repair function, alongside elevated cGAS-STING signaling and type I interferon (IFN-I) responses in human AD brains compared to non-AD. In the PS19 transgenic (PS19Tg) mouse model of tauopathy, we found significantly elevated levels of DDSBs and altered expression of DNA repair proteins during early stages of disease, which preceded the dysregulation of cGAS-STING signaling and emergence of significant neuropathology in the later stage. Interestingly, genetic deletion of cGAS shifted microglial polarization from a pro-inflammatory M1 phenotype toward an anti-inflammatory M2 state, accompanied by a reduction in IFN-I signaling and improved cognitive performance in PS19Tg mice. Pharmacological STING inhibition reshaped the transcriptomic landscape, revealing selective regulation of pathways governing synaptic plasticity, and immune responses. This transcriptional reprogramming was accompanied by suppression of inflammatory responses, reduction in synaptic pathology, and attenuation of tau pathology in PS19Tg mice, underscoring STING as a therapeutic target for tauopathy. In conclusion, our findings reveal that genotoxic-immune crosstalk drives neuroinflammation and tau pathology and identify a conserved, druggable cGAS-STING axis that can be targeted to impede or slow disease progression in tauopathies.

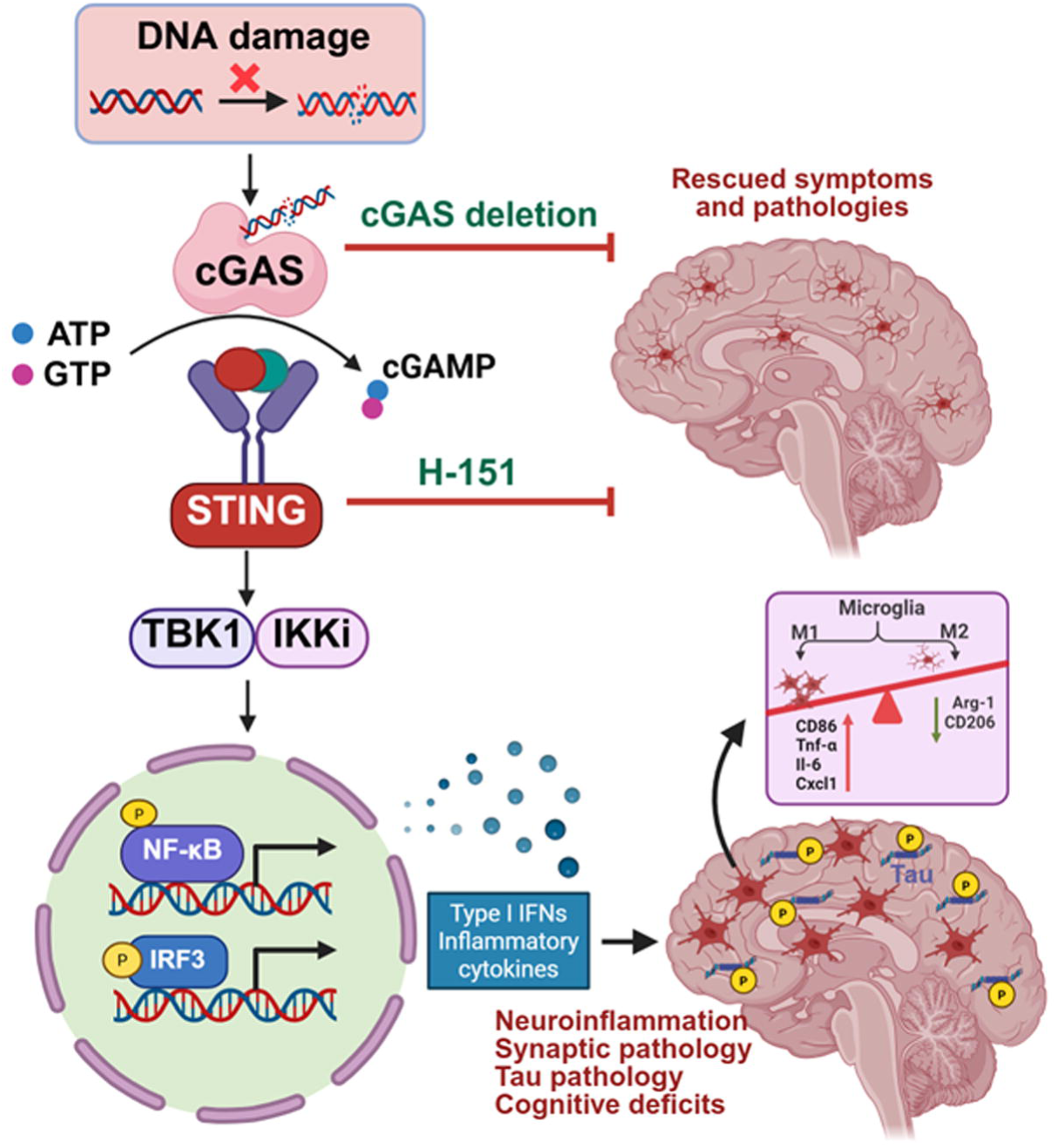

## Introduction

Tauopathies, including Alzheimer’s disease (AD) and frontotemporal dementia (FTD), are neurodegenerative disorders characterized by the abnormal buildup of hyperphosphorylated and misfolded tau in neurons and glial cells. This aberrant tau aggregation disrupts synaptic integrity and causes neuronal loss, ultimately leading to cognitive decline (Samudra et al., 2023, Wang et al., 2024). The global burden of dementia is projected to increase dramatically, reaching 131.5 million cases by 2050. Despite this concerning trend, the mechanisms behind tau pathology during aging are not well understood, and there are no disease-modifying treatments available. Elucidating the upstream molecular pathways that drive tau-mediated neurodegeneration is essential for uncovering novel therapeutic targets and advancing effective strategies to halt or reverse disease progression.

The human genome is continuously exposed to DNA damage, yet the brain relies on sophisticated repair mechanisms to preserve genomic stability. Among these, DNA double-strand breaks (DDSBs) represent the most severe form of damage, accumulating with age. Failure of DNA repair affects transcription and translation, resulting in breakdowns for genomic integrity and is associated with decreased expression of genes linked to neuronal function and plasticity (Delint-Ramirez and Madabhushi, 2025, Konopka and Atkin, 2022, Thadathil et al., 2019). Unresolved DDSBs have been observed in various neurological disorders (Delint-Ramirez and Madabhushi, 2025, El-Saadi et al., 2022, Madabhushi et al., 2014, Maynard et al., 2015, Neven et al., 2024, Thadathil et al., 2021). Moreover, an emerging body of evidence underscores the interplay between DNA damage and tauopathy (Cimini et al., 2022, Khurana et al., 2012, Li et al., 2024, Zheng et al., 2020). Interestingly, DNA-repair-deficient mice develop tau pathology in the brain (Hou et al., 2018), further supporting the link between DNA damage and tau pathology. Tau-expressing flies exhibited over a two-fold increase in comet tail length, indicating heightened DNA damage (Bukhari et al., 2024). The regulation of DNA repair mechanisms leads to improvements in health and cognitive function in a C. elegans model of tauopathy (Tiwari et al., 2024).

Despite the pathogenic role of DDSB in tauopathies, the molecular mechanism underlying DDSB-mediated neuropathology during tauopathy remains poorly understood. Recent studies have highlighted the critical interplay between DDSBs, and immune signaling in neurodegeneration, with a specific focus on the cyclic GMP-AMP synthase (cGAS)-stimulator of interferon genes (STING) pathway (Chung et al., 2024, Gulen et al., 2023, Hinkle et al., 2022, Khan et al., 2025). cGAS, a key sensor of cytosolic DNA, detects DDSB and catalyzes the synthesis of cyclic GMP-AMP (cGAMP), which activates the STING pathway (Chen et al., 2016). This activation leads to the recruitment of TANK-binding kinase 1 (TBK1), which phosphorylates key transcription factors, including interferon regulatory factor 3 (IRF3) and nuclear factor κB (NF-κB) (Balka et al., 2020, Huang et al., 2023, Paul et al., 2021, Zhang et al., 2024). These transcription factors drive type I interferon (IFN-I) responses and the production of pro-inflammatory cytokines, amplifying neuroinflammation (Chen et al., 2016, Zhang et al., 2024). Damaged nuclear DNA can also activate the non-canonical STING pathway, which operates independently of cGAS and primarily activates NF-κB (Dunphy et al., 2018). The cGAS-STING pathway regulates the generation and secretion of senescence-associated secretory phenotype (SASP) factors, including pro-inflammatory cytokines driven by IRF3 and NF-κB (Herbstein et al., 2024). Activation of the cGAS-STING pathway has been detected in postmortem human brains from individuals with Parkinson’s disease (Hinkle et al., 2022, Khan et al., 2025), Alzheimer’s disease (Xie et al., 2023), and amyotrophic lateral sclerosis (Marques et al., 2024), as well as in mouse models of tauopathy and amyloidosis (Chung et al., 2024, Jin et al., 2021, Swarup et al., 2019, Thanos et al., 2025, Udeochu et al., 2023, Xie et al., 2023, Yan et al., 2025). This evidence highlights the pivotal role of cGAS-STING in the neuroinflammatory processes that accelerate neurodegeneration, suggesting that modulating this pathway may hold therapeutic potential for tauopathies.

In this study, we demonstrated that accumulation of DDSBs combined with reduced DNA repair capacity, drives cGAS-STING-mediated inflammation in human AD postmortem brains. Using cell culture systems and mouse models, we show that early DDSB accumulation precedes tau pathology through dysregulation of the cGAS–STING pathway. Importantly, inhibition of cGAS mitigates neuroinflammation and cognitive deficits by promoting a protective shift in microglial polarization, while pharmacological blockade of STING with H-151 similarly suppresses neuroinflammatory responses and tau pathology. Overall, our results uncover a mechanistic link between genotoxic stress and innate immune activation in tauopathies, positioning the cGAS-STING pathway as a novel and actionable therapeutic target

## Materials and Methods

### Human post-mortem tissues

Postmortem human brain tissues of AD and cognitively normal (Non-AD) were acquired from the Michigan Brain Bank in Ann Arbor, Michigan. **Table S1** provides a summary of the clinical features of the patients and the controls. The University of Tennessee Health Science Center Institutional Review Board approved the study, which was carried out in accordance with accepted ethical standards (IRB # 20-07595-NHSR; Exempt Application 874552), in Memphis, Tennessee, USA. The handling of the human samples complied with all personal protection safety regulations. Formalin-fixed and frozen post-mortem brain tissues with the entorhinal cortex from AD and non-AD groups were examined for proteins involved in DNA damage and repair and immune responses.

### Histology of human AD and non-AD brains

The immunohistochemical examination was conducted on coronal sections at the entorhinal cortex level as described previously by us (Javed et al., 2020, Khan et al., 2025, Thadathil et al., 2021). Following deparaffination, sections were pretreated with 10 mM citric acid buffer (pH 6.0) for antigen retrieval. Sections were washed and blocked in 5% bovine serum albumin (BSA; Sigma Aldrich # A7906) for an hour. The sections were incubated with the primary antibodies for STING (1:200, Proteintech), and Iba1 (1:500; Synaptic System). Following washing, the sections were incubated for 1 hour with Alexa Fluor-labeled secondary antibodies mixed with DAPI solution (# 62248; ThermoFisher Scientific Inc), then mounted using ProLong™ Diamond Antifade Mountant (#P36961; ThermoFisher Scientific Inc). An investigator who was blinded to the groups examined 9 microscopic fields taken from 3 sections of each subject’s brain under 400X magnification to determine the number of STING positive microglia cells.

### Relative quantitative real-time reverse-transcriptase PCR (RT-qPCR)

Utilizing Roche’s LightCycler® 480 System and primers from the supplemental material, SYBR green-based RT-qPCR was used to compare the relative mRNA levels of DNA repair genes and cGAS-STING immune responses in human and mouse entorhinal cortex tissues (Table S2). Using random primers and following the instructions provided by ThermoFisher Scientific’s RETROscript™ Reverse Transcription Kit, cDNA was produced. GAPDH (human or mouse) was used as internal control. Relative gene expression was calculated using 2−^ΔΔCT^ method and expressed as fold change. The primers sequences are provided in supplementary **Table S2**.

### Mice and brain tissues

All mouse experiments were carried out in compliance with the Institutional Animal Care and Use Committee’s approval and followed the National Institutes of Health’s Guidelines for the Care and Use of Laboratory Animals. In-house bred PS19 transgenic mice (PS19Tg) were periodically back-crossed to the C57BL/6J wild-type line. In the PS19Tg mouse model, the prion protein promoter drives expression of the mouse T34 isoform of tau, carrying the human P301S mutation, with one N-terminal insert and four microtubule-binding repeats (1N4R). cGAS knockout mice (Jackson Laboratory, strain #026554) were crossed with PS19Tg mice to generate PS19Tg/cGAS^-/-^ offspring.

### Behavioral Assessments

*<u>Novel object recognition.</u>* The novel object recognition test (NOR) task is a behavioral assay used to assess recognition memory in rodents, based on their spontaneous preference for novel over familiar stimuli (Botton et al., 2010, Cruz-Sanchez et al., 2020). Testing was conducted in a blue acrylic chamber (40 × 40 × 30 cm), containing two objects. On the first day (habituation), mice were individually placed in the arena and allowed to explore freely for 10 minutes. To minimize olfactory interference, the chamber was cleaned with 75% ethanol between trials. On the second day (acquisition phase), animals were allowed to settle for 30 minutes in the behavior room before being presented with two identical red objects, placed 8 cm from the adjacent walls. After a 2-hour retention interval, one of the familiar objects was replaced with a novel object (a rectangular cuboid), while the other remained unchanged. Each test lasted 5 minutes, and mouse behavior was recorded using EthoVision XT 17 video tracking software (Noldus, Wageningen, The Netherlands), with particular attention to the time spent exploring the novel object. The recognition index (RI), calculated as RI = TN/(TN+TF), reflects the proportion of exploration directed toward the novel object and served as an indicator of working memory. TN is the time spent exploring the novel object, and TF is the time spent exploring the familiar object.

### Immunofluorescence staining

Immunofluorescence staining was carried out according to our prior descriptions (Khan et al., 2018, Khan et al., 2025, Thadathil et al., 2021). Under anesthesia, the brains of mice were swiftly removed, quickly placed in 4% paraformaldehyde solution, and immediately cryoprotected with 30% sucrose in 0.1M PBS at 4°C for 48-72 hr. Brain sections (20 µm) were cut on a cryostat, washed with PBS, and then blocked with 5% BSA and 0.3% Triton X-100. Sections were incubated overnight at 4°C with primary antibodies: Iba-1, STING, P-IRF3, CD86, Phospho-Tau (Ser202, Thr205), PSD95 or synaptophysin. Following washes with PBS, Alexa Fluor-labeled secondary antibodies (1:500, Invitrogen, USA) were applied to sections for 1 hr. Following washing, the sections were incubated for 1 hour with Alexa Fluor-labeled secondary antibodies mixed with DAPI solution. Afterward, the sections were washed and mounted using ProLong™ Diamond Antifade Mountant. An observer who was blind to genotype performed an average cell count using a fluorescent microscope at 400X magnifications using three distinct fields for each region of each mouse’s hippocampus and cortex. **Table S3** lists all antibodies used for immunohistochemistry and Western blot, along with their sources, catalog numbers, and working dilutions

### Immunoblot analyses

Hippocampal or cortical tissues from each human subject or mouse brain were isolated and homogenized in ice-cold RIPA buffer containing Halt™ protease and phosphatase inhibitor (ThermoFisher Scientific, USA). Supernatants were obtained by centrifuging the lysates at 12,000 rpm for 20 minutes. Proteins were separated on 4-20% Criterion™ TGX™ Precast Midi Protein Gel (Bio-Rad, USA) by SDS-PAGE and transferred into PVDF membranes using a wet transfer system. After blocking in 5% BSA for 1 hour, membranes were incubated overnight at 4°C with primary antibodies: γ-H2A.X (Ser139), cGAS, STING, Phospho IRF3, Phospho-TBK1, Phospho-Tau (Ser396), Phospho-Tau (Ser202, Thr205), PSD95, Arginase-1, CD206, NLRP3, GAPDH, or α-tubulin in tris-buffered saline with Tween 20 (TBST) containing 5% BSA. After washing with TBST, membranes were incubated with the HRP-conjugated secondary antibodies for 2 hours at room temperature with gentle rocking. Protein bands were visualized using an ECL detection kit (SuperSignal™ West Pico or Femto; Thermo-Fisher Scientific, USA). The Odyssey Fc imaging system (Li-Cor, USA) was used to capture images of the membranes. NIH ImageJ was used to quantify protein bands, with normalization to the corresponding loading control.

### RNA sequencing

RNA from hippocampal tissue was isolated and measured using Qubit 4 and Qiagen Qiaxcel. RNA libraries were prepared using the Illumina stranded RNA library kit. Libraries were sequenced on Illumina NextSeq 2000 using a P3 sequencing kit. All FASTQ files were obtained from the sequencer and quality control was performed using FASTQC. Reads with a Phred score below Q20 were trimmed. The cleaned reads were aligned to the mm38 reference genome using RNA STAR. Resulting SAM files were processed to extract gene-level read counts, which were then normalized across samples using the TMM method. Principal component analysis (PCA) and Pearson correlation plots were generated from the normalized transcriptome data. Differential expression analysis was conducted with DESeq2, and genes with a P-value ≥ 0.05 or fold change ≤ 1.5 were excluded. Benjamini-Hochberg correction was applied to control the false discovery rate, and genes with an FDR ≥ 0.05 were removed. The final list of significant differentially expressed genes was imported into R for heatmap visualization and further analyzed with Gene Set Enrichment Analysis (GSEA) and STRINGdb for pathway and gene ontology enrichment.

### Cell cultures and immunocytochemistry

Primary neurons were cultured from embryonic day 14 (E14) PS19Tg mouse embryos, as described previously (Guo and Lee, 2013, Khan et al., 2014, Thadathil et al., 2021). Cells were counted, plated in 24-well culture dishes with coverslips coated with Matrigel and poly-L-lysine, and grown in vitro for two weeks in Neurobasal A media with 2% B27, 1% FBS (ATCC # 30-2020), and 1% mM L glutamine. Culture media were replaced every three days, and cultures were kept at 37 °C in an incubator that was humidified with 5% CO2. The cultures (DIV 6) were exposed to 1.5 μg tau K18 PFFs and Aβ oligomers (derived from 7PA2 CHO cells) (Cleary et al., 2005, Podlisny et al., 1995) or vehicle to promote Aβ-tau aggregation and neurodegeneration (ATN), an essential feature of AD as described by us and others (Guo and Lee, 2013, Thadathil et al., 2021). The culture continued to grow for 21 days and was then fixed for immunostaining.

To assess DDSB, fixed cells were twice washed in 1X PBS, permeabilized for 15 minutes with 0.3% Triton X-100 and then blocked with 2.5 % BSA in PBS. Cells were then incubated for 2 hours at room temperature with γ-H2A.X (Ser139) and MAP2A/B primary antibodies, followed by washing and incubation for 1 hour with fluorescently labeled secondary antibodies in the presence of DAPI. After three additional PBS washes, cells were mounted on slides with coverslips for imaging and subsequent quantification.

### Statistical analysis

GraphPad Prism version 10.2 (GraphPad Software, San Diego, CA) was used for all statistical analyses. Comparisons between two groups were assessed with an unpaired two-tailed t-test, while one-way ANOVA followed by post-hoc testing was applied to examine treatment effects. Results are expressed as mean ± SEM, and differences were considered significant at P < 0.05.

## Results

### Genotoxic stress and dysregulated cGAS-STING immune responses in human AD brains

We previously reported elevated DDSB and altered DNA repair function in the hippocampus of AD brains (Thadathil et al., 2021). In the present study, we demonstrate that increased DDSBs cooperate with the cGAS– STING pathway to amplify IFN responses in the entorhinal cortex of AD brains. Specifically, we observed elevated levels of γ-H2A.X (Ser139), a marker of DDSBs, in AD brains compared to non-AD control (**Fig. 1 A, B**). To further investigate DNA repair deficits, we examined the expression of key DNA repair proteins, including MRE11, RAD50, BRCA1, and 53BP1. Notably, MRE11, RAD50 (**Fig. 1 E, F)**, and BRCA1 **(Fig. S1A**) expression were significantly reduced in human AD brains, suggesting impaired DNA damage recognition and repair. However, 53BP1 expression remained unchanged (**Fig. 1 G)**. Previous findings from our and others lab demonstrated cell-type-specific differences in 53BP1 expression, with distinct patterns in neurons and glia (Shanbhag et al., 2019, Thadathil et al., 2021). This suggests that while neurons may experience compromised DNA repair, glial cells could respond differently to genotoxic stress. The selective downregulation of MRE11, RAD50, and BRCA1 highlights potential defects in DDSB repair, which may exacerbate genomic instability and contribute to neurodegeneration in AD.

**Fig. 1.**
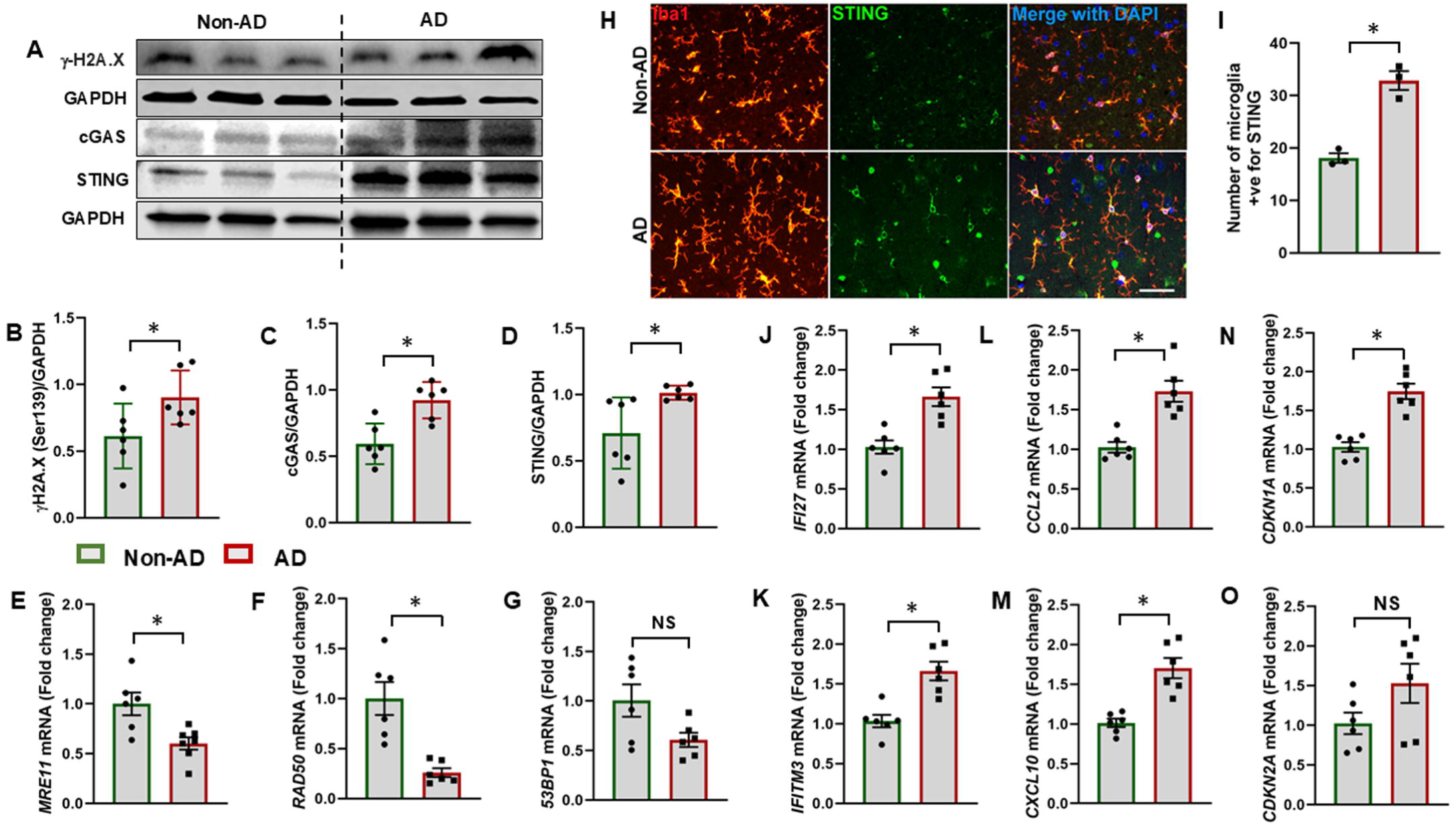
Increased genotoxic stress and cGAS-STING immune response in human AD brains. **(A-D).** Representative immunoblots of γ-H2A.X (Ser139), cGAS and STING proteins and their quantification in entorhinal cortex of AD and non-AD brains. (N=6/group). **(E-G).** The mRNA expression of DNA repair genes (*MRE11, RAD50,* and *53BP1*) assessed by RT-qPCR in entorhinal cortex of AD and non-AD brains. (N=6/group). **(H-I).** Representative fluorescent images of Iba1 (microglia; red) and STING (green) in entorhinal cortex of AD brains and their quantification. Scale bar = 50 µm (N=3/group). **(J-O).** RT-qPCR was used to analyze the mRNA expression of *IFN genes (IFI27, IFITM3), chemokines (CCL2 and CXCL10) and senescence (CDKN1A and CDKN2A)* in entorhinal cortex of AD and non-AD brains. Group differences were analyzed using a two-tailed t-test (N=6/group). Data are presented as mean ± SEM. P < 0.05; NS: not significant

DNA damage has been widely implicated in triggering innate immune responses through intracellular DNA sensors, particularly cGAS. In our analysis, we detected a significant increase in both mRNA (**Fig. S1B, C)** and protein expression levels of cGAS and STING in the entorhinal cortex of AD brains compared to non-AD controls (**Fig. 1 C, D**). Immunohistochemical analysis revealed significantly increased STING expression in microglial cells in human AD compared to non-AD (**Fig. 1 H, I**). This upregulation indicates enhanced activation of the cGAS-STING pathway in microglia, reflecting a heightened immune surveillance response to genotoxic stress in AD pathology, which may contribute to neuroinflammation. Activation of the cGAS-STING pathway was further linked to an amplified IFN-I responses, as demonstrated by the upregulation of IFN-response genes, including *IFI27* and *IFITM3* (**Fig. 1 J-K**). Additionally, we observed increased mRNA expression of the chemokines *CCL2* and *CXCL10* (**Fig. 1 L-M**), which are known to mediate microglial activation and neuroinflammatory cascades. Beyond its role in neuroinflammation, cGAS-STING activation is also implicated in cellular senescence, a state of irreversible growth arrest that has been increasingly recognized as a driver of aging and neurodegenerative diseases. In line with this, we observed upregulation of *CDKN1A* and *CDKN2A* mRNA expression in the entorhinal cortex of AD brains compared to non-AD (**Fig. 1 N-O**).

### Early genotoxic stress in the brains of PS19Tg mice

We first examined the accumulation of DDSB in neuronal cultures derived from PS19Tg mice, The cultures were treated with tau PFFs and Aβ oligomers, recapitulating amyloid, tau pathology and neurodegeneration (A/T/N) conditions. We found robust expression of γH2A.X (Ser139) in neuronal cells treated with tau PFFs and Aβ (**Fig. 2 A-B**). Next, we examined the expression of DDSBs in the hippocampus and cortex of 6-month-old PS19Tg mice. Immunoblot analyses revealed a significant accumulation of DDSBs, with markedly increased γ-H2A.X (Ser139) levels in hippocampus (**Fig. 2 C and D**) and cortex regions (**Fig. S2A and B**). Using RT-qPCR, we quantified the mRNA expressions of *Mre11, Rad50, and 53BP1* which are critical components of DNA damage sensing and repair pathways. We observed a significant downregulation of *Mre11,* and *Rad50*, suggesting impaired DNA repair (**Fig. 2 E and F**). However, mRNA expressions of 53BP1 remained unchanged (**Fig. 2 G**). These findings indicate that early impairments in DNA damage sensing and repair mechanisms drive the accumulation of DDSBs in PS19Tg mice, potentially amplifying genomic instability and neurodegeneration in tauopathies at later stage of disease.

**Fig. 2.**
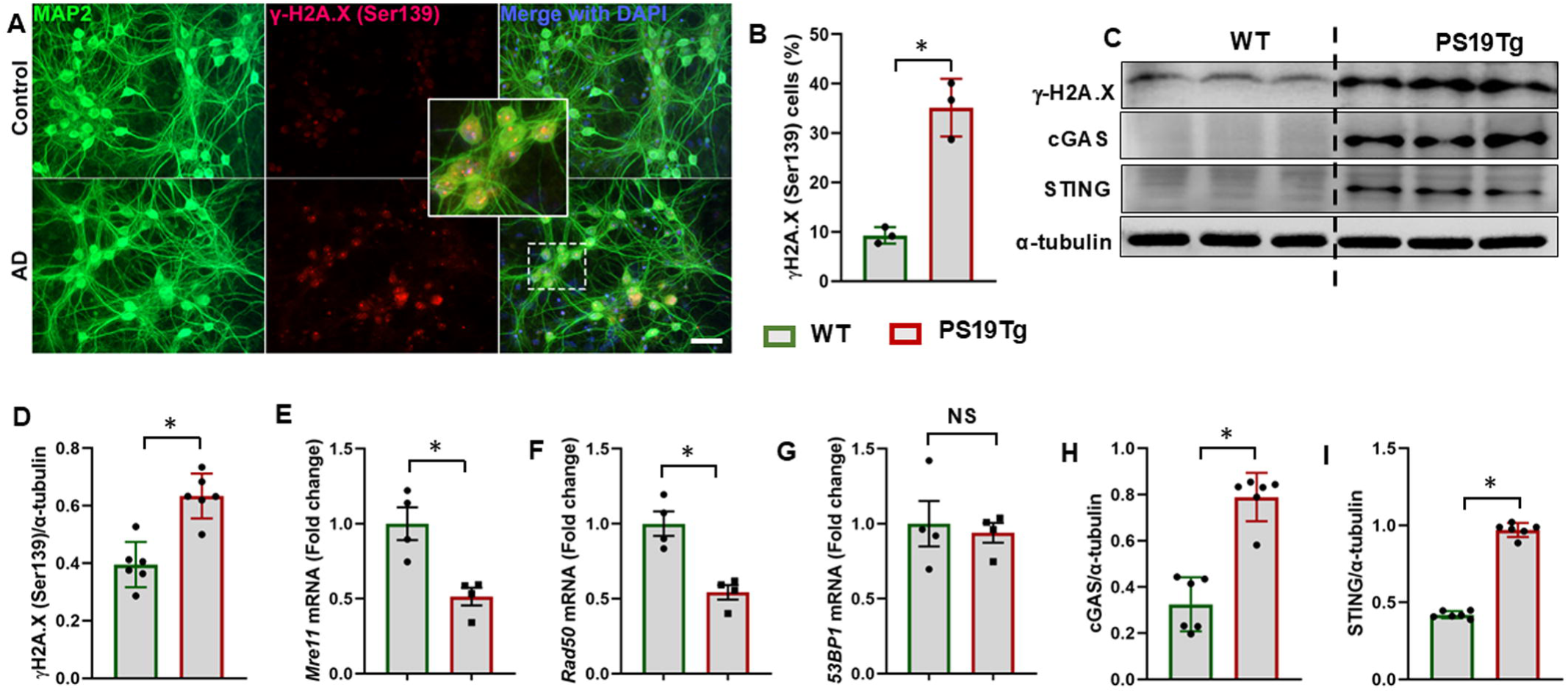
Neural genotoxicity and c-GAS-STING dysregulation in tauopathy. **(A and B).** Primary neuronal cultures derived from PS19Tg mice were exposed with Tau PFF, Aβ oligomers or vehicle. Representative immunofluorescence images show increased DDSB expression (γ-H2A.X (Ser139); red) in neurons (MAP2; green) following treatment with Tau PFF and Aβ oligomers compared with vehicle-treated cultures. Data are expresed as means ± SEM. Scale bar = 50 μm. **(C-G)**. Representative immunoblot analysis of γ-H2A.X (Ser139), and mRNA expression of DNA repair proteins (*Mre11, Rad50,* and *53BP1)* in the hippocampus of 6-month-old WT and PS19Tg mice (N=4/group). **(H and I).** Representative immunoblots analysis of cGAS and STING in the hippocampus of 6-month-old WT and PS19Tg mice (N=6/group). Data were compared between groups using a two-tailed t-test and represented as mean ± SEM *P < 0.05

### Temporal and spatial dysregulation of cGAS-STING signaling in PS19Tg mice

To assess the role of cGAS-STING signaling in tauopathy progression, we examined its dysregulation in the hippocampus and cortex of 6- and 9-month-old PS19Tg mice. Immunoblot analyses revealed a significant increase in cGAS and STING expression in hippocampus (**Fig. 2 C, H and I)** and cortex (**Fig. S2A, C and D**), early in the disease course (6 months). This upregulation was further exacerbated at 9 months in the hippocampus (**Fig. 3 A-C)** and cortex (**Fig. 3 F-H**) coinciding with worsening tau pathology (**Fig. S3A**). Consistent with these findings, immunofluorescence staining demonstrated a striking overexpression of STING and its downstream target phosphorylated IRF3 (p-IRF3) in activated microglia in the hippocampus (**Fig. 3 D and E**) and cortex (**Fig. 3 I-J**) of PS19Tg mice, compared to age-matched controls. This suggests that microglial activation in tauopathy is closely linked to cGAS-STING signaling, potentially driving chronic neuroinflammation.

**Fig. 3.**
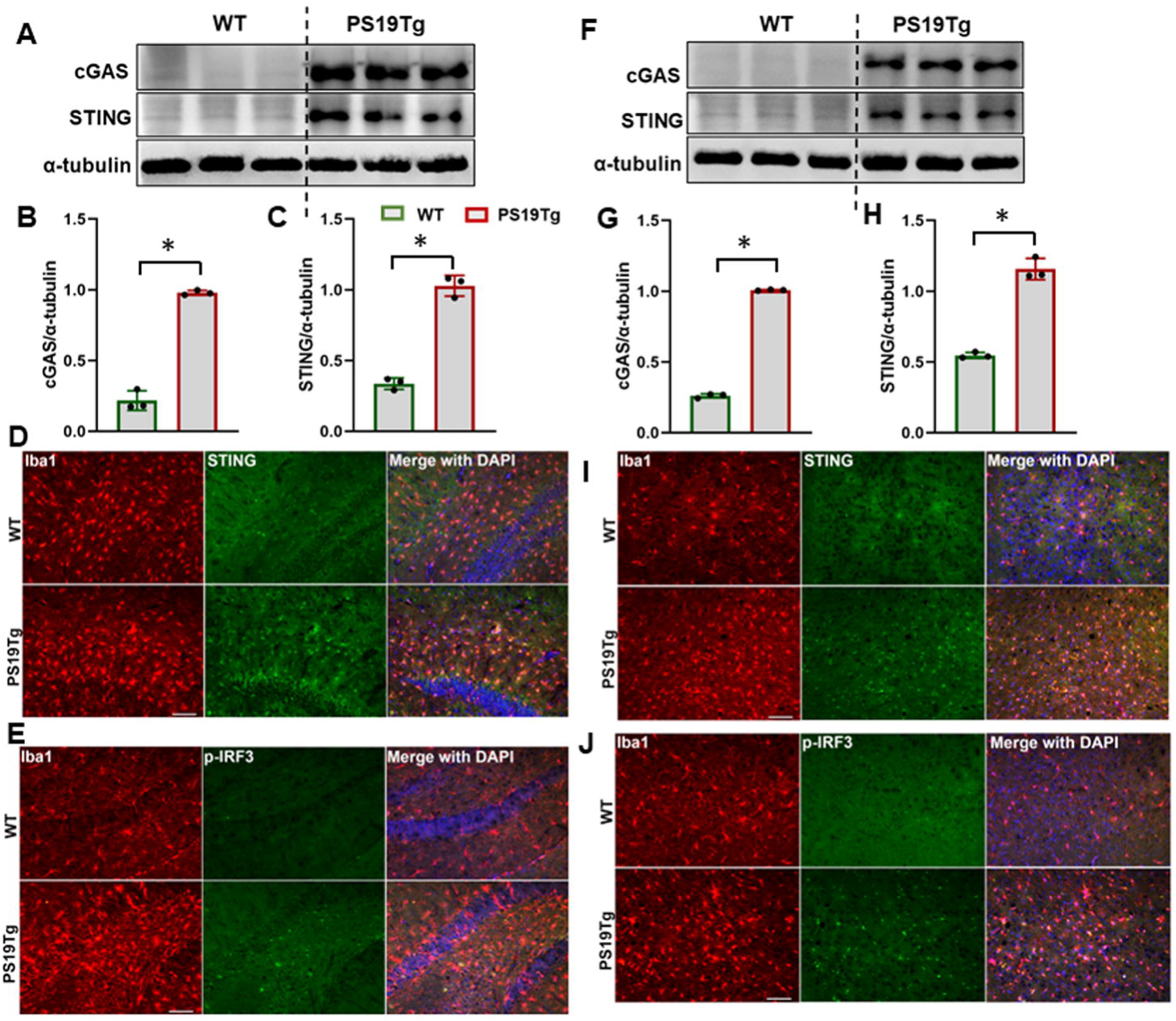
Induction of cGAS-STING-mediated interferon responses in tauopathy with significant disease. **(A-C).** Immunoblot of cGAS, and STING in the hippocampus of 9-month-old WT and PS19Tg mice and their quantification (N=3/group). **(D)** Representative immunofluorescence images of microglia [Iba1; red], STING (green) and DAPI (blue) **(E)** Representative immunofluorescence images of microglia [Iba1; red], p-IRF3 (green) and DAPI (blue) in the hippocampus of WT and PS19Tg mice and their quantification. Scale bar = 100 µm **(F-H).** Immunoblot of cGAS, and STING in the cortex of 9-month-old WT and PS19Tg mice and their quantification (N=3/group). **(I)** Representative immunofluorescence images of microglia (Iba1; red), STING (green) and DAPI (blue) **(J)** Representative immunofluorescence images of microglia (Iba1; red), p-IRF3 (green) and DAPI (blue) in the cortex of 9-month-old WT and PS19Tg mice and their quantification. Scale bar = 100 µm.

### cGAS deletion alleviates tau pathology and cognitive deficits through STING-IFN modulation

To investigate the contribution of the cGAS-STING pathway to neuroinflammatory responses in tauopathy, we analyzed brains from 8-month-old cGAS-deficient PS19Tg mice (PS19Tg/cGAS^-/-^) and PS19Tg (PS19Tg/cGAS^+/+^) mice. Western blot analyses revealed significantly reduced levels of STING, p-IRF3, and tau phosphorylation (PHF-13) in PS19Tg/cGAS^-/-^ mice compared to age-matched PS19Tg controls (**Fig. 4 A-D**). RT-qPCR confirmed efficient loss of cGAS transcript in these mice, validating the genetic knockout (**Fig. 4 E)**. Consistent with blunted STING signaling, we observed marked downregulation of IFN-stimulated genes, including *Isg15, Ifi27* and *Ifitm3*, which are canonical downstream targets of IRF3 (**Fig. 4 F-H)**. Furthermore, mRNA expressions of Sting and senescence marker *Cdkn2a* were significantly reduced (**Fig. S4 A-B)**, suggesting that cGAS loss disrupts the DNA damage-induced senescence program commonly observed in tauopathy. To evaluate the contribution of cGAS deficiency to cognitive impairment and tauopathy, we performed the novel object recognition (NOR) test in PS19Tg mice with or without cGAS. PS19Tg/cGAS^-/-^ mice showed a marked improvement in NOR performance, with a significantly increased recognition index (F (2, 21) = 9.892; P=0.0009) comparable to PS19Tg mice (**Fig. 4 I**). These results suggest that dysregulated cGAS contributes to tau-induced cognitive decline and that its loss confers protection against cognitive deficits in this tauopathy mouse model.

**Fig. 4.**
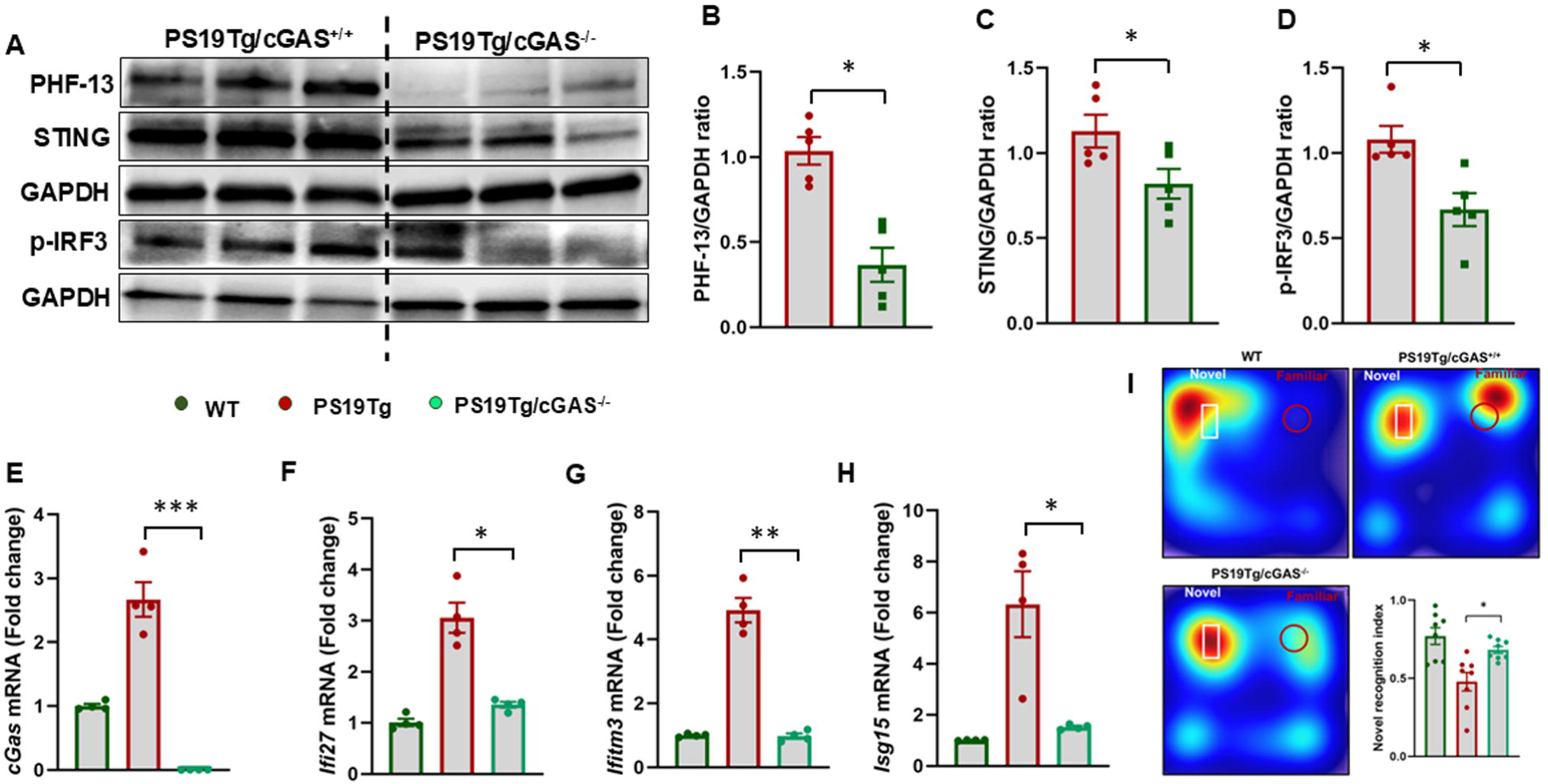
cGAS deficiency attenuates STING-mediated interferon response, reduces tau pathology and improves cognitive function. **(A-D).** Representative immunoblots of Phospho-Tau (PHF-13), STING, and p-IRF-3, and their quantification in the hippocampus of PS19Tg and cGAS deficient PS19Tg (PS19Tg/cGAS^-/-^) mice. *P < 0.05 (N=5/group). **(E-H).** RT-qPCR was employed to assess the mRNA expressions of cGas and interferon-stimulated genes (*Ifi27, Ifitm3* and *Isg15)* in the hippocampus of PS19Tg and cGAS-deficient PS19Tg (PS19Tg/cGAS^-/-^) mice. *P < 0.05 (N=4/group). **(I).** Recognition memory was assessed using the Novel Object Recognition (NOR) test. cGAS deficiency significantly improved recognition memory performance in PS19Tg mice. *P < 0.05 (N=8/group). Data were analyzed using a two-tailed t-test for comparisons between two groups or one-way ANOVA followed by Holm-Sidak multiple comparisons test and are presented as mean ± SEM *P < 0.05.

### cGAS deficiency alters microglial polarization toward an anti-inflammatory phenotype

We next assessed microglial polarization downstream of STING and IFN-I signaling to determine how cGAS-STING-mediated IFN responses influence microglial activation. To determine whether cGAS deficiency influences microglial polarization in tauopathy, we examined the expression of key pro- and anti-inflammatory markers in the brains of PS19Tg/cGAS^-/-^ mice (**Fig. 5**). Deletion of cGAS resulted in a pronounced shift in the microglial transcriptional profile toward an anti-inflammatory phenotype, with significant upregulation of M2-associated genes, including Arg1 and CD206, in PS19Tg mice (**Fig. 5 A-D**). In contrast, expressions of classical M1-associated pro-inflammatory genes, including *Cxcl1, Il6,* and *Tnf-α* were significantly reduced in PS19Tg/cGAS^-/-^ mice compared to PS19Tg mice (**Fig. 5 E-G)**. Immunohistochemistry analysis demonstrated a marked reduction in CD86 expression in microglia from cGAS-deficient PS19Tg mice compared to PS19Tg controls (**Fig. 5 H, I)**, indicating decreased microglial activation in the absence of cGAS signaling. CD86 is a well-established co-stimulatory molecule expressed by activated microglia, while Cxcl1, Tnf-α and Il6 are potent mediators of immune cell recruitment and cytokine-driven inflammation. The downregulation of these markers indicates diminished classical microglial activation. This polarization shift suggests that the absence of cGAS suppresses pro-inflammatory signaling and promotes a reparative, anti-inflammatory microglial state in tauopathy.

**Fig. 5.**
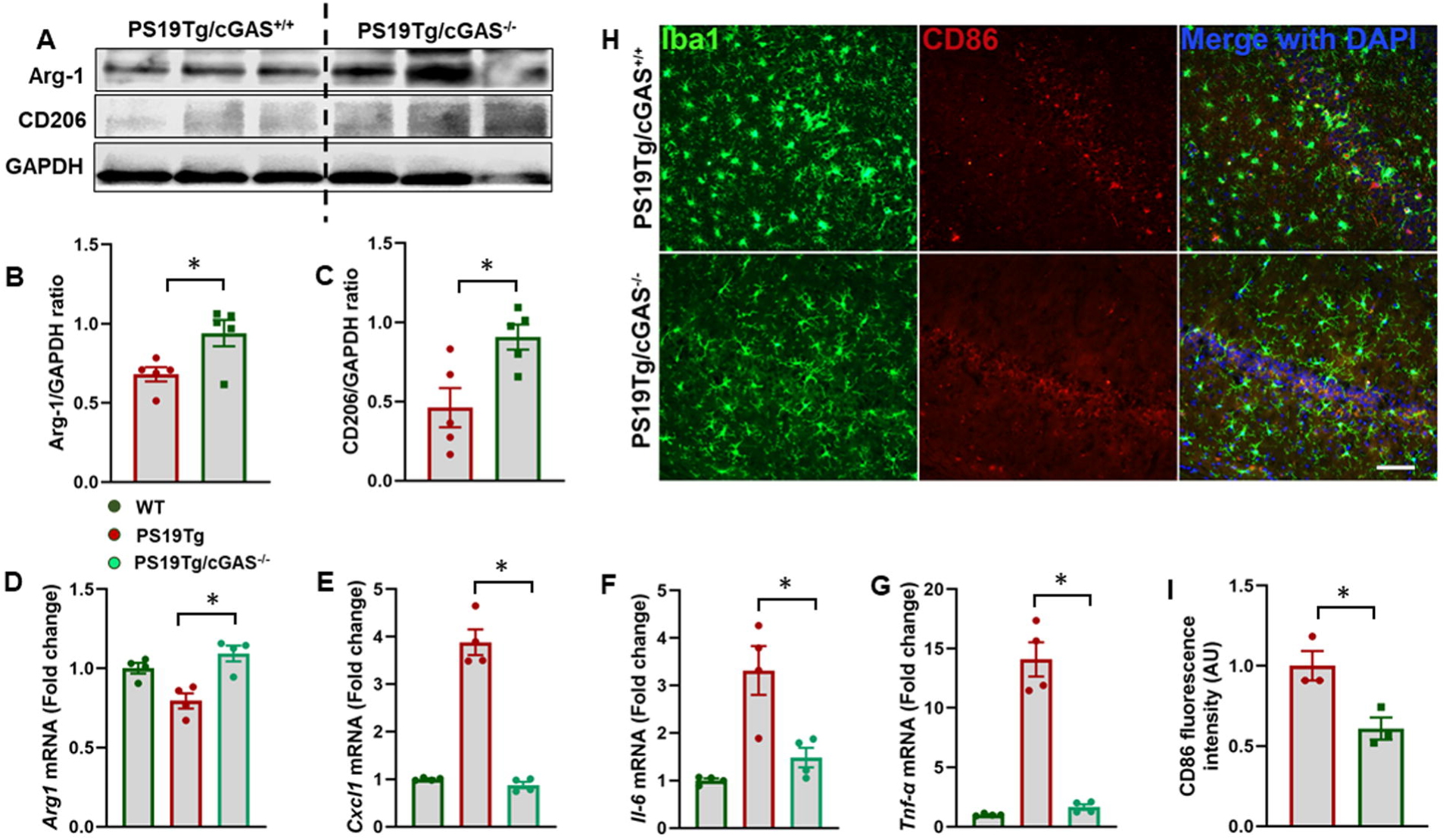
cGAS deficiency alters microglial polarization toward an anti-inflammatory phenotype. **(A-C).** Representative immunoblots of Arginase-1 (Arg-1) and CD206, and their quantification in the hippocampus of PS19Tg (PS19Tg/cGAS^+/+^) and cGAS-deficient (PS19Tg/cGAS^-/-^) mice. *P < 0.05 (N=5/group). **(D-G).** RT-qPCR was used to analyze the mRNA expression of M1 and M2 markers: *Arg-1, Cxcl1, Il-6,* and *TNF-α* in the hippocampus of PS19Tg/cGAS^+/+^ and PS19Tg/cGAS^-/-^ mice (N=4/group). **(H and I).** Representative immunofluorescence images of microglia (Iba1; green), CD86 (red) and DAPI (blue) in the hippocampus of PS19Tg/cGAS^+/+^ and PS19Tg/cGAS^-/-^ mice and their quantification (N=3/group). Scale bar = 100 μm. Data were analyzed using a two-tailed t-test for comparisons between two groups or one-way ANOVA followed by Holm-Sidak multiple comparisons test and are presented as mean ± SEM *P < 0.05.

### Pharmacological inhibition of STING alleviates IFN responses in PS19Tg mice

To determine whether pharmacological inhibition of STING could mitigate neuroinflammation and genotoxic stress in tauopathy, we treated 6-month-old PS19Tg mice with H-151, a selective and potent STING inhibitor, twice per week for 8 weeks (10 mg/kg, i.p.). H-151 administration has been previously demonstrated to suppress STING-IFN activation in neurological disorders (Khan et al., 2025, Xie et al., 2023, Yan et al., 2025). Our findings revealed that H-151 treatment effectively suppressed STING activation in the hippocampus of PS19Tg mice. Immunoblot analyses showed a significant reduction in STING, and its downstream target phosphorylated TBK1 (**Fig. 6 A-C**). Additionally, we found significant reduction in expression levels of IFN-I response genes *Ifi27, Ifitm3, Ifn-γ,* and *Isg15* (**Fig. 6 D-G**), and *chemokines Ccl2 and Cxcl10* (**Fig. 6 H, I**) in PS19Tg mice treated with H-151, indicating inhibition of STING-dependent inflammatory cascade. Furthermore, STING inhibition reduces senescence and SASP in PS19Tg mice, as evidenced by a significant reduction in the expression levels of *Il-6, Il-1β, TNF-α, Cdkn1a, and Cdkn2a,* (**Fig. 6 J-N**), suggesting that blocking STING signaling can alleviate senescence-associated neuroinflammatory responses in tauopathy.

**Fig. 6.**
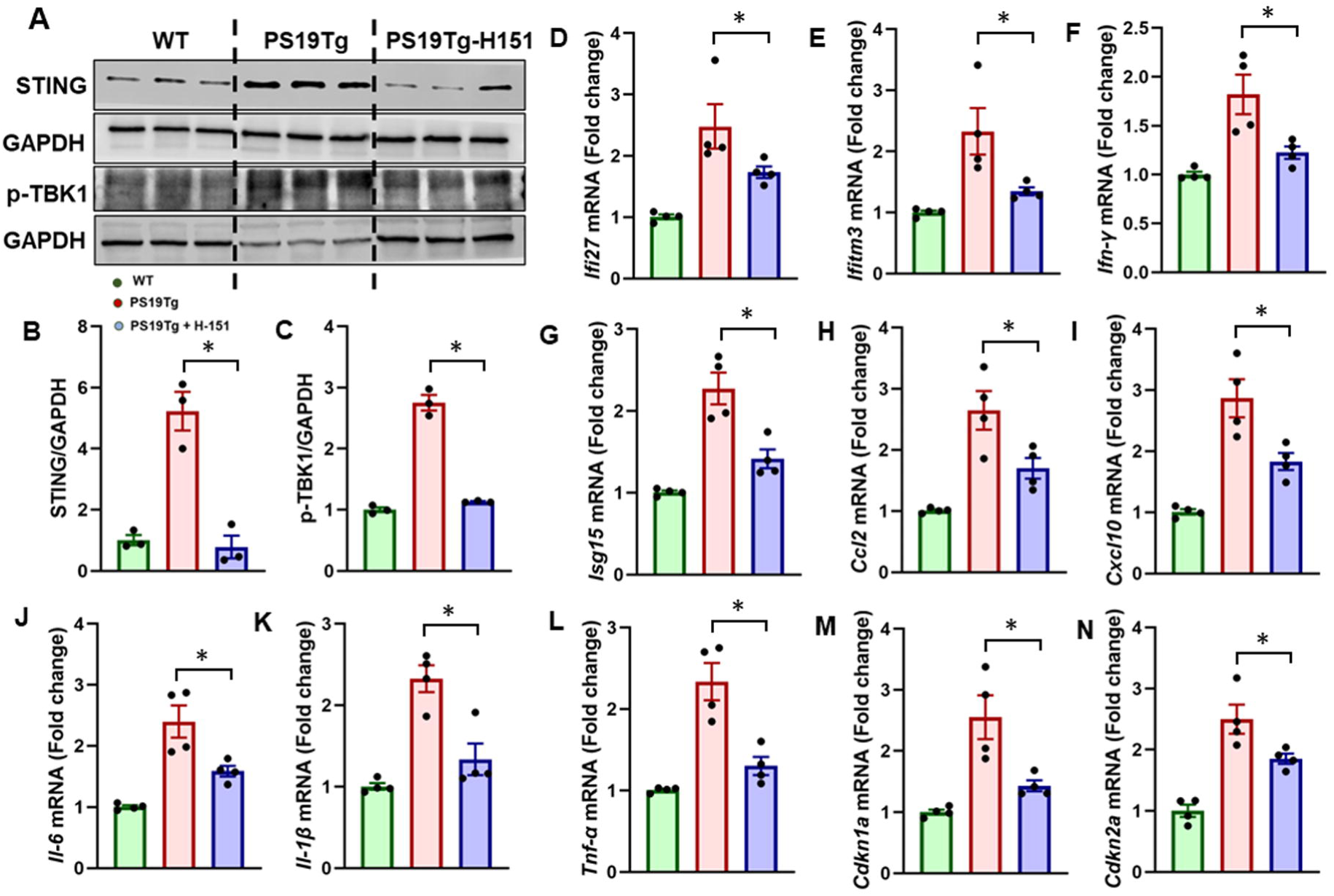
STING inhibition reduces IFN-I signaling and SASP gene expression in tauopathy. **(A-C).** Immunoblot of STING, and p-TBK1 and their quantification in the hippocampus of PS19Tg and H-151 treated PS19Tg mice. The values are expressed as mean ± SEM *P < 0.05 (N=3/group). **(D-N).** RT-qPCR was used to determine the mRNA expressions of interferon-stimulated genes (*Ifi27, Ifitm3, Ifn-γ, Isg15),* and SASP genes (*Ccl2,* Cxcl10, *Tnf-α, Il-6*, *Il-1β*, *Cdkn1a,* and *Cdkn2a*) in the hippocampus of PS19Tg and H-151 treated PS19Tg mice. Data are presented as mean ± SEM (N=4/group). Statistical analysis was performed using one-way ANOVA followed by the Holm-Sidak multiple comparisons test. Differences were considered significant at P < 0.05.

### STING modulation shapes transcriptomic profiles and reduces a broad range of tau pathology

RNA-sequencing analysis revealed 2,385 differentially-expressed genes (DEGs) following treatment with the H-151 in the PS19Tg mouse model (**Fig. 7 A-C**). Among these, 514 genes were upregulated, indicating potential activation of protective or reparative processes and 1,871 genes were downregulated, consistent with suppression of disease-associated pathways. To gain insight into the biological relevance of these transcriptional changes, we performed Gene Set Enrichment Analysis (GSEA), which identified 100 significantly enriched pathways (FDR < 0.05). These included pathways involved in inflammatory signaling, cognition/memory, immune responses, IFN response, and synaptic plasticity. A total of 195 genes contributed to the enrichment of these 100 pathways, suggesting a focused regulatory shift affecting key functional networks. In parallel, STRING-based Gene Ontology (GO) analysis was conducted to further annotate and contextualize the DEGs. This approach revealed 541 significantly enriched GO terms, spanning biological processes, molecular functions, and cellular components. These findings suggest that the H151 not only broadly reshapes the transcriptome but also converges on specific molecular pathways associated with synaptic plasticity, neuroinflammatory responses, and DNA damage responses, highlighting potential mechanisms of action and therapeutic relevance.

**Fig. 7.**
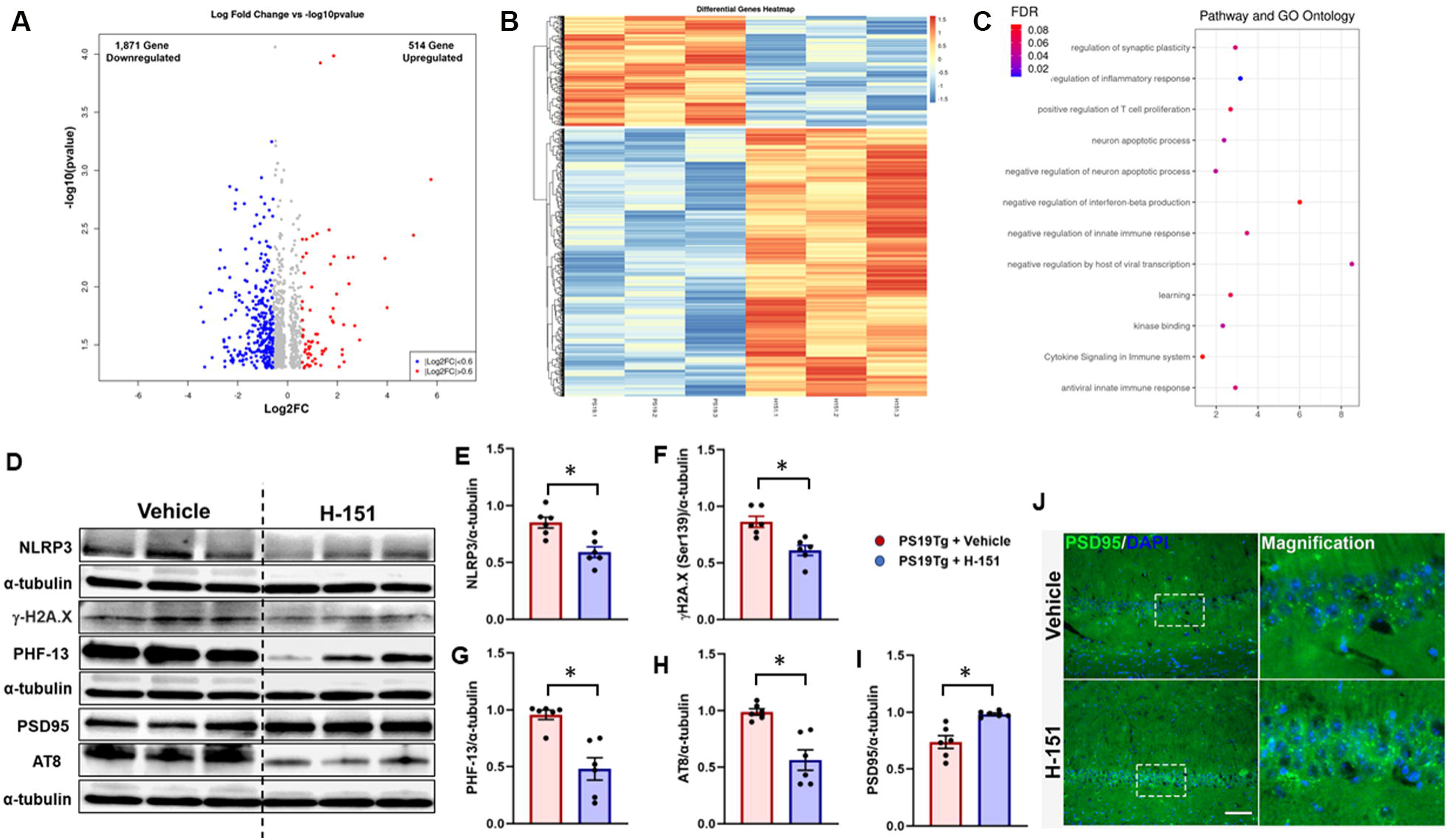
Transcriptomic profiling reveals pathways modulated by STING inhibition. (A) Volcano plot showing differentially-expressed genes (DEG) between H-151 treated PS19Tg mice and PS19Tg controls. Red dots represent genes with significant upregulation, and blue dots represent genes with significant downregulation (B) Heat map of DEGs across samples, highlighting clusters of genes altered by H-151 treatment. (C) Pathway and Gene Ontology (GO) enrichment analyses of DEGs, showing key signaling pathways and biological processes regulated by H-151 treatment (N=3/group). **(D-I).** Representative immunoblots of NLRP3, γ-H2A.X, PHF-13 (pS396), PSD95, and AT8 (pS202, t205) and their quantification in the hippocampus of 8-month-old PS19Tg and H-151 treated PS19Tg mice. The values are expressed as mean ± SEM *P < 0.05 (N=6/group). **(J).** Representative immunofluorescence images of hippocampal sections stained for PSD95 in PS19Tg mice treated with vehicle or H-151. DAPI (blue) was used to visualize nuclei. Scale bar: 100 µm. (N=3/group).

Further molecular analyses demonstrated that pharmacological blockade of STING produced broad beneficial effects across multiple pathological domains in PS19Tg mice. Immunoblot assessments showed significantly reduced expression of NLRP3 expression (**Fig. 7 D, E**), indicating dampened inflammasome activation. Strikingly, DNA damage markers were significantly reduced following treatment (**Fig. 7 F**), suggesting that STING inhibition interrupts the deleterious feedback loop between neuroinflammation and genotoxic stress. Importantly, pathological tau phosphorylation was markedly decreased at key residues, including Ser396, and Ser202/Thr205, which are known to drive tau aggregation and neurotoxicity (**Fig. 7 G, H**). We further found that treatment with H-151 increased PSD95 expression (**Fig. 7 I, J**), a synaptic marker associated with cognitive integrity, suggesting that cGAS-STING inhibition not only mitigates neuroinflammation but also preserves synaptic function. Collectively, these results demonstrate that targeting the cGAS-STING pathway can simultaneously attenuate multiple hallmarks of tauopathy, including DNA damage, tau pathology, synaptic dysfunction, and inflammatory signaling. This work highlights the potential of STING inhibition as a therapeutic strategy to enhance neuronal and synaptic resilience in tau-driven neurodegenerative disease.

## Discussion

Neurodegenerative disorders, including tauopathies, are often characterized by aberrant immune responses that contribute to disease progression. Thus, there is significant interest in elucidating the upstream molecular pathways, such as the innate immune sensors and nucleic acid-sensing mechanisms that initiate and sustain these inflammatory cascades in tauopathy. In this study, we uncover that the early-stage genotoxic stress triggers cGAS-STING-mediated immune responses, driving aberrant tau pathology and cognitive deficits. Strikingly, genetic deletion of cGAS in the PS19Tg tauopathy model not only dampens STING-mediated IFN-I responses but also restrains pro-inflammatory microglial polarization, thereby mitigating cognitive deficits and tau pathology. Furthermore, pharmacological inhibition of STING reduces IFN-I-driven inflammation, ameliorates synaptic pathology, and diminishes aberrant tau accumulation. RNA-seq analysis reveals that STING inhibition reshapes the disease-associated transcriptome, particularly affecting pathways related to synaptic plasticity, immune activation, and DNA damage responses. By linking genotoxic stress to immune activation, our study uncovers a critical upstream driver of tau pathology and opens new avenues for disease-modifying interventions.

DNA damage accumulation is a well-documented feature of aging and various age-related neurological disorders (Delint-Ramirez and Madabhushi, 2025, Madabhushi et al., 2014, Thadathil et al., 2021, Wang et al., 2021). Previous research has highlighted the interplay between DNA damage and tau pathology (Asada-Utsugi et al., 2022, Cimini et al., 2022, Farmer et al., 2020, Khurana et al., 2012, Rossi et al., 2018, Sultan et al., 2011, Tiwari et al., 2024). For instance, tau-expressing Drosophila models exhibit increased comet tail length, reflecting enhanced DNA damage (Bukhari et al., 2024, Tiwari et al., 2024). Tau oligomers are also implicated in disrupting DNA repair mechanisms, exacerbating the accumulation of unresolved DNA damage and fueling neurodegeneration (Violet et al., 2015, Zheng et al., 2020). A study by Asada-Utsugi and coworkers have shown the coexistence of DDSBs and phosphorylated tau in the cortex of human AD brains (Asada-Utsugi et al., 2022). Despite the pathogenic accumulation of DDSBs in tauopathies, the molecular mechanisms driving tauopathy remain poorly understood. In human AD brains, we detected a significant accumulation of DDSBs, as indicated by elevated γ-H2AX (Ser139) levels along with decreased mRNA expression of *MRE11, RAD50,* and *BRCA1*. The MRE11-RAD50-NBS1 (MRN) complex, which is essential for DDSB detection and repair through HR and NHEJ pathways (Lamarche et al., 2010), showed a marked reduction in their expressions in both human AD brains and the tauopathy mouse model. This disruption in MRN function may significantly contribute to genotoxic stress, which likely accelerates disease progression in tauopathies. Interestingly, despite the downregulation of MRN complex components, the expression of 53BP1, a crucial DDR protein involved in the response to DNA damage, remained unchanged. This observation raises the possibility that 53BP1 may function in a cell-specific manner, with differential expression patterns in various cell types, particularly neurons and microglia. Previous studies, including our own, have demonstrated that 53BP1 is expressed differently in neurons and glia, suggesting distinct roles in these cells during DNA damage responses (Shanbhag et al., 2019, Thadathil et al., 2021). Consistent with our findings in human AD data, we observed an increased DDSB, assessed by γ-H2AX (Ser139), in the hippocampus and cortex of PS19Tg mice at 6 months of age. Moreover, we found a reduction in the expression of *MRE11* and *Rad51*. In contrast, 53BP1, a key player in the NHEJ pathway, remained unchanged. These findings indicate that although the NHEJ pathway is active, it may not function optimally in the context of tauopathy-induced genomic instability. These findings further support the notion that DNA damage, combined with impaired repair mechanisms, contributes to the pathophysiology of tauopathies, highlighting the importance of maintaining efficient DDR and repair systems in neurons.

DNA damage in tauopathies extends beyond genomic instability, influencing key disease-driving mechanisms such as neuroinflammation and neurodegeneration. One of the key pathways activated in response to unresolved DNA damage is the cGAS-STING signaling pathway, which has garnered increasing attention in neurodegenerative diseases, including tauopathies (Carling et al., 2024, Jin et al., 2021, Udeochu et al., 2023, Yan et al., 2025). Upon recognition of DDSBs, cGAS synthesizes cGAMP, which activates STING, a central mediator of the innate immune response. This activation triggers downstream signaling cascades, including the phosphorylation of TBK1 and IRF3, which in turn promotes the production of IFN-I and other pro-inflammatory cytokines. Chronic activation of this pathway leads to sustained neuroinflammation, a hallmark of neurodegenerative diseases (Gulen et al., 2023, Zhang et al., 2024). IFN-1 responses have been shown to inhibit DNA repair proteins, impairing the recruitment of DNA repair factors to sites of DNA damage (Ka et al., 2021). This deficit in DNA repair can lead to the accumulation of DNA breaks, which further activates cytosolic DNA-sensing pathways and amplifies IFN production, thereby closing a detrimental feedback loop that exacerbates neuroinflammation and drives tau pathology. In addition to nuclear DNA damage, cGAS also senses mitochondrial DNA that leaks into the cytosol either through mitochondrial disruption or defective mitophagy (Quan et al., 2025). In alignment with these findings, our study reveals a significant dysregulation of the cGAS-STING pathway in human AD brains. Our findings further revealed increased expression of IFN genes in AD brains, including *IFI27*, *IFITM3, CCL2, CXCL10,* and senescence-related genes *CDKN1A* and *CDKN2A*. These results suggest an amplification of the neuroinflammatory response and the presence of cellular senescence in AD, which may contribute significantly to disease progression. CCL2 and CXCL10 are chemokines involved in recruiting immune cells, including microglia, to brain inflammation sites. Their upregulation supports the role of neuroinflammation, mediated by activated microglia and immune cells in AD. Increased chemokine expression in our data aligns with previous studies linking sustained neuroinflammation to neuronal damage and cognitive decline in tauopathies (Udeochu et al., 2023, Wojcieszak et al., 2022). We also observed elevated levels of *IFI27* and *IFITM3*, interferon-induced genes typically upregulated in response to viral infections and cellular stress. Their increased expression in AD brains suggests a chronic IFN-I response, likely driven by cGAS-STING pathway activation. This response has been implicated in amplifying neuroinflammation and worsening neurodegeneration in several diseases including tauopathy (Sanford et al., 2024, Udeochu et al., 2023). The upregulation of senescence markers *CDKN1A* and *CDKN2A*, possibly in response to DNA damage, highlights the presence of cellular senescence in AD brains. Senescent cells contribute to a pro-inflammatory state via senescence-associated secretory phenotype (SASP), which can exacerbate neurodegeneration, promote tau pathology, and accelerate cognitive decline (Ahmad et al., 2024, Gaikwad et al., 2024). Our findings provide compelling evidence that increased DNA damage, dysregulated immune responses, and cellular senescence are key drivers of neuroinflammation in tauopathies, reinforcing the cGAS-STING axis as a potential therapeutic target.

In the PS19Tg tauopathy model, temporal and spatial analyses at 6 and 9 months revealed pronounced activation of the cGAS-STING pathway, evidenced by elevated cGAS, STING, and phosphorylated IRF3 in both the hippocampus and cortex, highlighting early and region-specific innate immune engagement in tauopathy. In our study, we observed significant activation of microglia expressing key markers of cGAS-STING pathway activation, including phosphorylated IRF3. This activation indicates that microglia are involved in the innate immune response triggered by DNA-sensing cGAS-STING pathway in tauopathies. The sustained activation of the cGAS-STING pathway in microglia drives chronic neuroinflammation, which accelerates tau aggregation and neurodegeneration. This finding is consistent with previous studies linking chronic cGAS-STING activation to persistent neuroinflammation in neurodegenerative diseases, including tauopathies (Gulen et al., 2023, Udeochu et al., 2023, Xie et al., 2023, Yan et al., 2025). Taken together, the interplay between cGAS-STING signaling, and persistent DNA damage provides a mechanistic link between genomic instability and immune response in tauopathies.

Next, we investigated whether modulation of the cGAS-STING pathway could confer protection against neuroinflammation, cognitive decline, and tau pathology. Remarkably, cGAS deletion drove a marked shift from a pro-inflammatory, disease-associated microglial state toward a more homeostatic transcriptional profile, revealing a pivotal role for cGAS-STING signaling in shaping microglial responses in tauopathy. This polarization shift was accompanied by a reduction in IFN-I signaling and expression of key inflammatory mediators, suggesting that cGAS functions upstream of microglial activation pathways in tau-driven neuroinflammation. Given the growing evidence that microglial states critically shape neurodegenerative outcomes, our data position cGAS as a pivotal regulator of innate immune reprogramming in the diseased brain. These findings are consistent with prior studies implicating nucleic acid sensors in the modulation of microglial activity (Jiang et al., 2021, Kong et al., 2024) and extend them by identifying a functional role for cGAS in shaping the neuroinflammatory landscape in tauopathy. Beyond modulating microglial polarization, cGAS deficiency also suppresses key inflammatory and neurodegenerative pathways. In PS19Tg mice, loss of cGAS markedly reduced STING activation and downstream effectors, including phosphorylated IRF3 and IFN-I responses, with concomitant suppression of interferon-stimulated genes. This dampening of innate immune signaling was associated with decreased tau phosphorylation and improved cognitive performance. Our findings are consistent with previous published studies identifying cGAS-STING signaling as a key mediator of innate immune activation in neurodegenerative diseases (Xie et al., 2023, Yan et al., 2025). These findings suggest that cGAS is an upstream amplifier of STING-mediated neuroinflammation, linking cytosolic nucleic acid sensing to both tau pathology and functional decline.

STING activation has been shown to drive NLRP3 inflammasome activation, promoting proinflammatory cytokine production and contributing to synaptic degeneration (Chung et al., 2024, Zhang et al., 2022). In line with prior studies demonstrating that STING inhibition alleviates AD-related pathology (Chung et al., 2024, Thanos et al., 2025, Xie et al., 2023), our findings show that treatment with H-151 effectively suppressed STING-IFN signaling and downstream NLRP3 activation. The concurrent reduction in SASP gene expression indicates that STING inhibition mitigates cellular aging processes that contribute to tau-mediated neurodegeneration. These observations align with previous studies showing that cGAS-STING inhibition alleviates neuroinflammation, dampens microglial reactivity, and enhances cognitive resilience in models of neurodegenerative disease (Chung et al., 2024, Udeochu et al., 2023, Xie et al., 2023, Yan et al., 2025). Beyond suppressing inflammation, H-151 acts as a transcriptomic recalibrator, dampening neuroinflammatory and DNA damage response pathways while restoring gene networks essential for synaptic integrity. Rather than merely silencing inflammation, H-151 appears to rewire the degenerative program, shifting cells toward a more resilient, pro-synaptic state. The decrease in phosphorylated tau (p-tau) levels following STING inhibition is particularly noteworthy, as tau hyperphosphorylation is a key driver of synapse loss and cognitive deficits in tauopathies. Pathological tau species are frequently localized to dendritic spines, where they disrupt synaptic structure and function, amplifying neuronal vulnerability. In our study, H-151 treatment not only reduced p-tau accumulation but also restored synaptic marker PSD95 and synaptophysin, indicating that dampening STING signaling can directly preserve synaptic integrity. This effect is likely mediated, at least in part, through suppression of NLRP3 inflammasome activation and downstream proinflammatory signaling, which are known to exacerbate tau-mediated synaptic damage. Together, these findings suggest that STING inhibition interrupts the deleterious interplay between tau hyperphosphorylation, neuroinflammation, and synaptic degeneration, highlighting its potential as a therapeutic strategy to protect synapses and maintain neuronal function in tauopathies. In conclusion, our studies suggest that the crosstalk between DNA damage and STING signaling plays a critical role in cognitive decline and neuroinflammation in tauopathy. cGAS-STING inhibition not only reduces genotoxic stress and neuroinflammation but also attenuates pathological tau burden and preserves synaptic and cognitive function. These findings highlight the cGAS-STING pathway as a key driver of neuropathology in tauopathy and underscore the potential of targeting both DNA repair mechanisms and STING signaling as a promising therapeutic strategy for tauopathies and related disorders.

## Acknowledgements

This work was supported by the NIH grant R03AG075597, Alzheimer’s Association Award AARG-NTF-22-972518 and Department of Defense Award Number HT9425-23-1-0043.

## Data availability

The data generated and analyzed in this study are available from the corresponding author upon reasonable request.

